# Choice of birth place among antenatal clinic attendees in rural mission hospitals in Ebonyi State, South-East Nigeria

**DOI:** 10.1101/520213

**Authors:** Leonard O. Ajah, Fidelis A. Onu, Oliver C. Ogbuinya, Monique I. Ajah, Benjamin C. Ozumba, Anthony T. Agbata, Robinson C. Onoh, Kenneth C. Ekwedigwe

**Affiliations:** Department of Obstetrics and Gynaecology, Faculty of Medical Sciences,University of Nigeria, Ituku-Ozalla Campus, Enugu; Department of Obstetrics and Gynaecology, Federal Teaching Hospital, Abakaliki; Institute of Maternal and Childhealth,University of Nigeria, Ituku-Ozalla Campus, Enugu

**Keywords:** Birth Place, Antenatal Clinic Attendees, Rural, Nigeria

## Abstract

**Background:** Low utilization of health facilities for delivery by pregnant women poses a public health challenge in Nigeria.

**Aim:** To determine the factors that influence the choice of birth place among antenatal clinic attendees.

**Methodology:** This was a cross-sectional study of the eligible antenatal clinic attendees at Mater Misericordiae Hospital, Afikpo and Saint Vincent Hospital, Ndubia in Ebonyi State from February 1, 2016 to June 30, 2016. Analysis was done using EPI Info 7.21 software (CDC Atlanta Georgia).

**Results:** A total of 397(99.3%) completely filled questionnaires were collated and analysed. Approximately 71% of the health facilities closest to the respondents had maternity services. It took at least 1 hour for 80.9% of the respondents to access health facilities with maternity services. Most (60.2%) of the respondents had antenatal care attendance and majority of them did so at public hospitals. Approximately 43.8% of the respondents were delivered by the skilled birth attendants. The common determinants of birth place were nearness of the health facilities, familiarity of healthcare providers, improved services, sudden labour onset and cost. Also 61.7% of the respondents chose to deliver in public health facilities due to favourable reasons but this could be hampered by the rudeness of some healthcare providers at such facilities. A significant proportion of private health facilities had unskilled manpower and shortage of drugs.

**Conclusion:** A greater proportion of women will prefer to deliver in health facilities. However there are barriers to utilization of these facilities hence the need for reversal of this ugly trend.

## Introduction

According to the United Nations International Children’s Emergency Fund (UNICEF), a woman dies from complications of childbirth every minute. Therefore about 529,000 women die annually from complications of childbirth.^1^ A woman in sub-Saharan Africa has 1 in 16 chances of dying in pregnancy or childbirth, compared to 1 in 4,000 risks in developed countries.^1^ This makes maternal mortality the health indicator showing the greatest disparity between the developing and developed countries.^2^ The 2013 Nigerian Demographic and Health Survey showed that the estimated maternal mortality ratio during the seven-year period prior to the survey was 576 maternal deaths per 100,000 live births while infant mortality rate was 69 deaths per 1,000 live births.^3^ The under-five mortality rate was 37 per 1000 live births while the neonatal mortality rate was 31 per 1000 births.^3^Antenatal clinic attendance in Nigeria is about 61% and only 38% of the deliveries are carried out by skilled birth attendants.^3^ A significant proportion of mothers in developing countries still deliver at home unattended to by skilled birth attendants.^4^ Individual factors that influence choice of birth place include maternal age, parity, education, marital status, household factors including family size and household wealth. Others are community factors such as socioeconomic status, community health infrastructure, region, rural/urban residence, availability of the health facilities and distance to such health facilities.^5^

To achieve the sustainable Development Goal 3, it is required that at least 80% of all deliveries should take place in a health facility.^6^ To enhance the utilization of health facilities during delivery in the country, barriers/determinants for utilizing such facilities during antenatal care and delivery among women need to be identified across all geographical regions.^7–9^ Previous studies have shown that sub-optimal care caused by lack of medical personnel, medical equipment, unavailability of drugs, high patient load and delay in attending to patients in the health facility, cultural beliefs and the influence of decision makers are some of the reasons patients choose to be delivered by the traditional birth attendants.^8,9,11–15^ Other factors which encourage home delivery are poverty, aversion to caesarean section, distance and means of transport to the health facility and separation of the patients in labour from their family members at the health facility.^10–14^

All pregnancies are at risk, and complications during pregnancy, delivery and the postpartum period are difficult to predict. In developing effective strategies for increasing the utilization of health facility delivery, it is necessary to understand the factors affecting choice of birth place. The underlying cause of low health facility-based delivery especially in rural areas needs further investigation and exploration in order to be better understood by reproductive health planners. This informed the need to determine and analyze the factors that influenced pregnant women’s choice of place of delivery at Mater Misericordiae Hospital, Afikpo and Saint Vincent Hospital, Ndubia, two rural mission hospitals in Ebonyi State, Nigeria. This study was aimed at determining the factors that influence choice of birth place among antenatal clinic attendees at rural Mission hospitals in Ebonyi State.

## Materials and methods

### Study sites

Mater Misericordiae Hospital,Afikpo was founded in 1946 by the St Patrick Missionaries, a Roman Catholic based religious group. It is a secondary hospital that serves Afikpo community, the neighboring communities in Ebonyi and other surrounding States of Abia, Akwa Ibom, Cross River, Enugu and Imo. The bulk of patients treated in this hospital are low income earners and rural dwellers. Saint Vincent’s Hospital,Ndubia is also a secondary hospital located in Izzi Local Government Area of Ebonyi State. The hospital was established in the early 1960 by the Catholic missionaries. The hospital serves the rural population in Izzi, Ikwo, Ezza and neighbouring states of Cross River and Benue. Bulk of the patients are rural dwellers and are predominantly farmers.

### Study design

This was a cross-sectional study in which interviewer-administered semistructured questionnaires were used to extract information from the consenting antenatal clinic attendees who had history of previous delivery prior to the index pregnancy. The questionnaires were administered by the researchers and trained research assistants. Prior to this study, the questionnaires were pre-tested on 20 antenatal clinic attendees at the Federal Teaching Hospital,Abakaliki. Each of the questionnaires had six sections and these sections comprised the participants’ socio-demographic characteristics, accessibility to healthcare services, previous antenatal and delivery history, rating of public and private health facilities, the pregnancy risks and recommendation for safer obstetric career. The number of study participants recruited from each centre was based on the proportion of antenatal clinic attendees in each of the study centres. The study participants were consecutively recruited until the number allocated to each centre was completed. All the consenting antenatal clinic attendees who had history of delivery within 3 years prior to the index pregnancy, irrespective of the outcome, were eligible to participate in the study. However nulliparous women, pregnant women who had prior history of delivery for more than 3 years before the index pregnancy and those who, despite adequate counselling, declined to participate in the study, were excluded. The minimum sample size for the study was calculated using the sample estimation formula for cross sectional studies,^15^

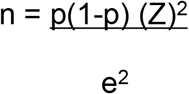

Where: n = sample size; p = prevalence of delivery in Nigerian health facilities = 38%^3^;1-p = number of deliveries outside established health facility =1-0.38=0.62 e = standard error = 5% and Z= standard normal variance =1.96 at 95% confidence interval. Adding a 10% attrition rate, n was 398.

Data were analyzed with EPI Info 7.21 software (CDC Atlanta Georgia). The ethical clearance for this study was obtained from the Ethics Committee of the Federal Teaching Hospital, Abakaliki. Institutional Permission was also obtained from Mater Misericordiae Hospital, Afikpo and St. Vincent Hospital,Ndubia before the commencement of this study. Confidentiality was ensured on all the study participants.

## Results

A total of 250(62.5%) and 150(37.5%) questionnaires were distributed at Mater Misericordiae Hospital, Afikpo and Saint Vincent’s Hospital,Ndubia respectively. However,397(99.3%) questionnaires were completely filled. Therefore the information from 397 questionnaires was collated and analyzed. Table 1 shows the socio-demographic characteristics of the participants. Majority of the participants were between 25 and 34 years, Christians, grand multiparous, married, Igbos, farmers and had primary level of education. Majority of the respondents’ husbands had secondary level of education and were farmers. Table 2 shows the respondents’ accessibility to healthcare services. It took majority of the study participants more than 5 hours to access the healthcare facility and most of the healthcare facilities are privately owned. Though 71% of the health facilities closest to the respondents had maternity services, 29% of such facilities did not have maternity services. It took at least 1 hour for 80.9% of the respondents to access health facility with maternity services and majority (34%) of them access such facilities by trekking. The respondents’ previous antenatal history is shown in table 3. Majority (60.2%) of the respondents had antenatal care attendance in their previous pregnancy and most of the previous ANC attendees did so at public hospitals and for more than 4 times. Table 4 shows the respondents’ determinants of birth place at the last delivery. Majority (43.8%) of the respondents were delivered by the skilled birth attendants in their last delivery. The most common determinant for the choice of birth place by the respondents was nearness of the health facility to the respondents. Majority (57.2%) of the respondents did not have transport fare to take them to health facility with maternity services. Majority (42.8%) of the respondents took the decision on birth place themselves. Table 5 shows the respondents’ choice of birthplace at the index pregnancy. Majority (61.7%) of the respondents chose to deliver in public health facilities and the main reasons for such choices were cost, better services and familiarity of the health workers to the respondents. The respondents’ rating of services in the health facilities is shown in table 6. The respondents answered that majority of health workers in public health facilities rendered good services, were skilled and had good infrastructure and enough drugs. However a significant proportion of the public health workers were said to be rude to patients. The participants also responded that majority of the health workers in private health facilities were caring, render good services and had good quality of infrastructure but had shortage of drugs. However, the participants responded that a significant proportion of private health workers were unskilled.

**Table 1:** The socio-demographic characteristics of the respondents.

| Age | Frequency | % |
| --- | --- | --- |
| 15 – 24 | 87 | 21.92 |
| 25 – 34 | 256 | 64.48 |
| 35 – 44 | 54 | 13.60 |
| **Religion** |  |  |
| Christianity | 347 | 87.41 |
| Islam | 20 | 5.04 |
| African traditional religion | 30 | 7.55 |
| **Parity** |  |  |
| 1 | 100 | 25.19 |
| 2-4 | 138 | 34.76 |
| ≥ 5 | 159 | 40.05 |
| **Ethnicity** |  |  |
| Igbo | 357 | 89.92 |
| Hausa | 14 | 3.53 |
| *Others | 26 | 6.55 |
| **Marital status** |  |  |
| Married | 304 | 76.57 |
| Single | 34 | 8.56 |
| Windowed | 33 | 8.32 |
| Divorced/separated | 26 | 6.55 |
| **Level of education** |  |  |
| No formal education | 76 | 19.14 |
| Primary | 149 | 37.53 |
| Secondary | 104 | 26.20 |
| Tertiary | 68 | 17.13 |
| **Husband educational level** |  |  |
| No formal education | 92 | 23.17 |
| Primary | 52 | 13.10 |
| Secondary | 176 | 44.33 |
| Tertiary | 77 | 19.40 |
| **Occupation of the respondent** |  |  |
| House wife | 76 | 19.14 |
| Farming | 169 | 42.57 |
| trading | 90 | 22.67 |
| Civil/public service | 62 | 15.62 |
| **Husband occupation** |  |  |
| Farming | 172 | 43.32 |
| Trading | 117 | 29.47 |
| Civil/public service | 108 | 27.21 |

**Table 2:** Respondents’ accessibility to healthcare services.

| **Time it took to reach the nearest health facility from home** | **Frequency (N=397)** | **Percentage** |
| --- | --- | --- |
| Close(<1hour) | 109 | 27.46 |
| Far (1-4hours) | 99 | 24.94 |
| Very far (≥5hour) | 189 | 47.60 |
| **Type of nearest health facility** |  |  |
| Public | 177 | 44.58 |
| Private | 220 | 55.42 |
| **Availability of maternity services in nearest health facility** |  |  |
| Yes | 282 | 71.03 |
| No | 115 | 28.97 |
| **Closest maternity services where nearest health facility does not have maternity services.** | **N=115** |  |
| Close(<1hour) | 22 | 19.13 |
| Far (1-4hours) | 58 | 50.43 |
| Very far(≥5hours) | 35 | 30.44 |
| **Means of transportation to health facility with maternity services** |  |  |
| Foot | 136 | 34.26 |
| Bicycle | 44 | 11.08 |
| Motorcycle | 90 | 22.67 |
| Vehicle | 127 | 31.99 |

**Table 3:**
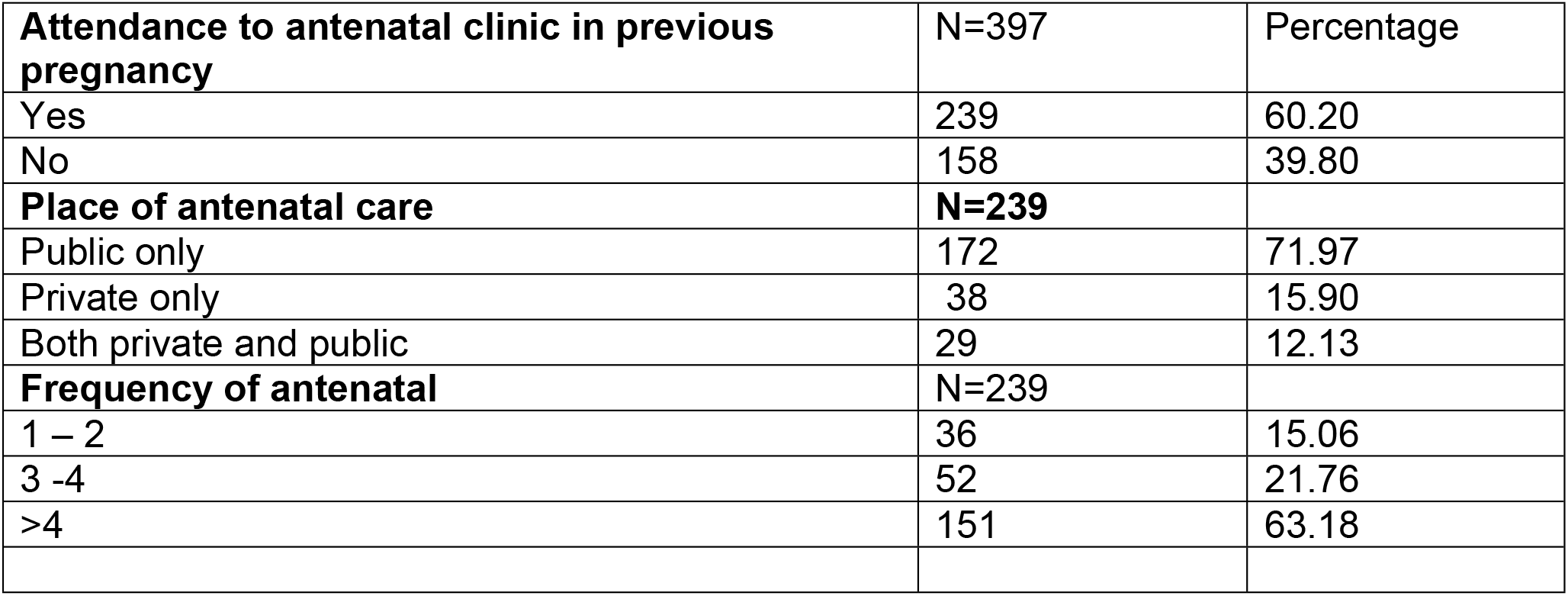
The respondents’ previous antenatal history.

|  |  |  |
| --- | --- | --- |
| **Attendance to antenatal clinic in previous pregnancy** | **N=397** | **Percentage** |
| Yes | 239 | 60.20 |
| No | 158 | 39.80 |
| **Place of antenatal care** | **N=239** |  |
| Public only | 172 | 71.97 |
| Private only | 38 | 15.90 |
| Both private and public | 29 | 12.13 |
| **Frequency of antenatal** | **N=239** |  |
| 1 – 2 | 36 | 15.06 |
| 3 -4 | 52 | 21.76 |
| >4 | 151 | 63.18 |

**Table 4:**
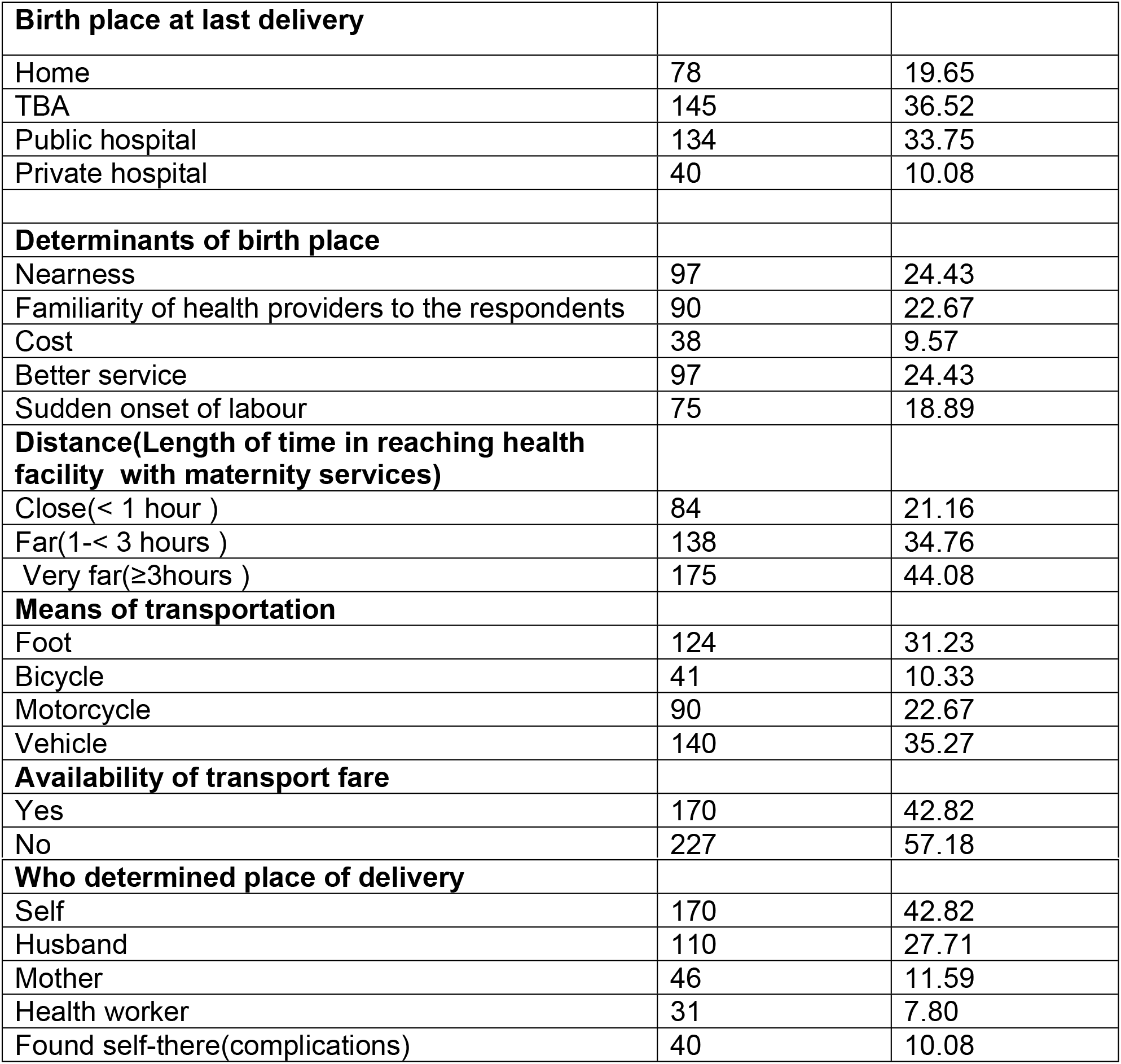
The respondents’ determinants of birth place at the last delivery.

**Table 5:**
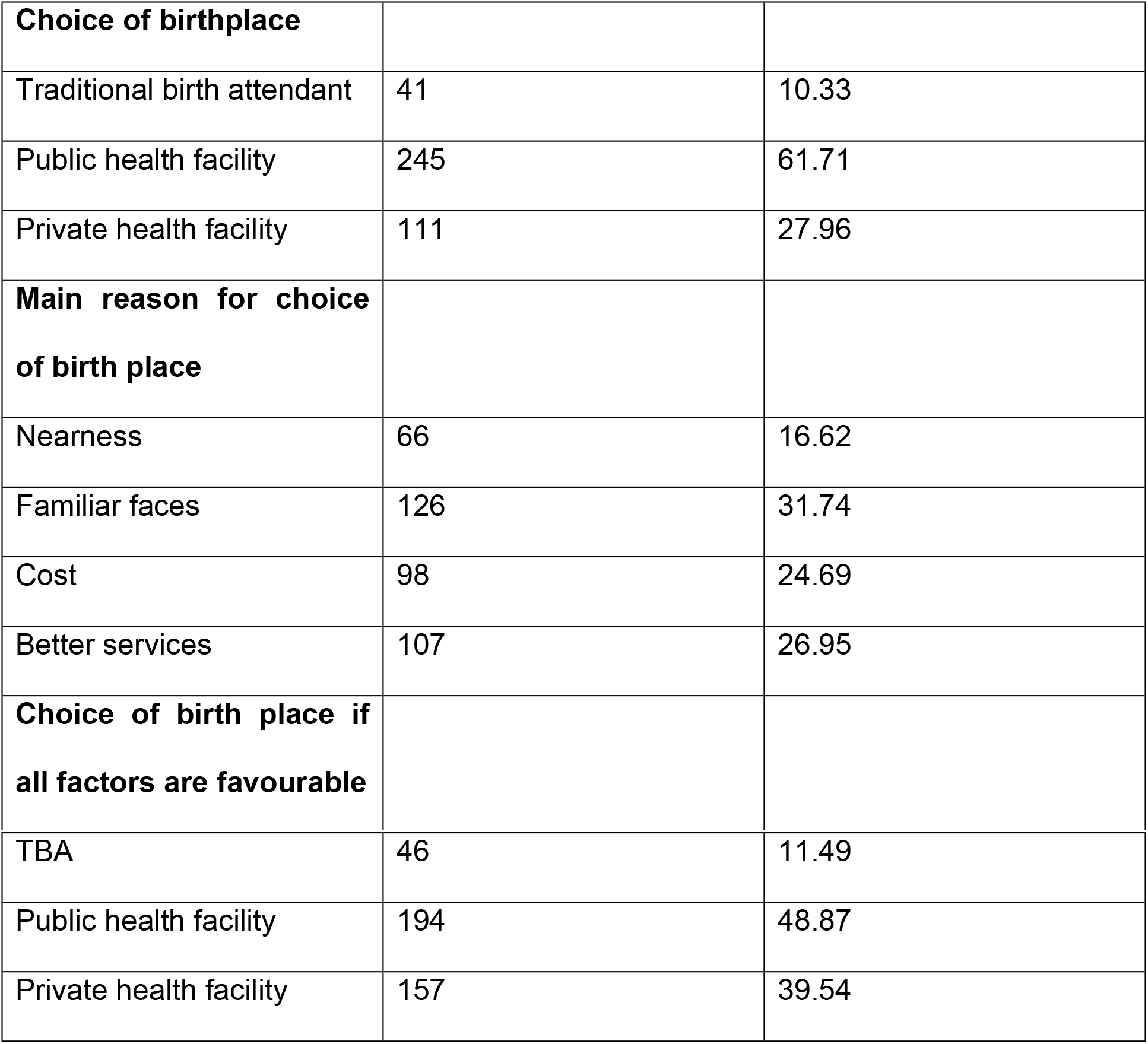
The respondents’ choice of birthplace in the index pregnancy.

|  |  |  |
| --- | --- | --- |
| **Choice of birthplace** |  |  |
| Traditional birth attendant | 41 | 10.33 |
| Public health facility | 245 | 61.71 |
| Private health facility | 111 | 27.96 |
| **Main reason for choice of birth place** |  |  |
| Nearness | 66 | 16.62 |
| Familiar faces | 126 | 31.74 |
| Cost | 98 | 24.69 |
| Better services | 107 | 26.95 |
| **Choice of birth place if all factors are favourable** |  |  |
| TBA | 46 | 11.49 |
| Public health facility | 194 | 48.87 |
| Private health facility | 157 | 39.54 |

**Table 6:** The respondents’ rating of services in the health facilities.

| Input | Indicator | Public |  | Private |  |
| --- | --- | --- | --- | --- | --- |
|  |  | No | % | No | % |
| *Health workers attitude | Rudeness | 269 | 46.30 | 78 | 13.43 |
|  | Caring | 185 | 31.84 | 350 | 60.24 |
|  | Sympathetic | 127 | 21.86 | 153 | 26.33 |
| Skills | Skilled | 321 | 80.86 | 151 | 38.04 |
|  | Unskilled | 76 | 19.14 | 246 | 61.96 |
| Quality of services | Excellent | 88 | 22.17 | 38 | 9.57 |
|  | Good | 175 | 44.08 | 200 | 50.38 |
|  | Fair | 101 | 25.44 | 22 | 5.54 |
|  | Poor | 33 | 8.31 | 137 | 34.51 |
| Total |  | 397 | 100 | 397 | 100 |
| Quality of infrastructure | Excellent | 116 | 29.22 | 53 | 13.35 |
|  | Good | 174 | 43.83 | 181 | 45.59 |
|  | Fair | 79 | 19.90 | 122 | 30.73 |
|  | Poor | 28 | 7.05 | 41 | 10.33 |
| Total |  | 397 | 100 | 397 | 100 |
| Availability of medicine and supplies | Available | 307 | 77.33 | 111 | 27.96 |
|  | Not available | 90 | 22.67 | 286 | 72.04 |
| Total |  | 397 | 100 | 397 | 100 |

\*multiple answers were allowed.

## Discussion

This study showed that the healthcare facilities were far to most of the respondents in their previous deliveries. The bulk of such facilities were privately owned. Though 60.2% of the respondents attended antenatal care clinics in their previous pregnancies, It was only 43.8% of them that were delivered by skilled birth attendants. The most common determinant for the choice of birth place by the respondents was nearness to the health facility to the respondents. It took at least 1 hour for 78.9% of the respondents to reach health facility with maternity services and a significant proportion of them usually reached the health facility by vehicles. However most of the respondents could hardly afford the transport fare to take them to such facilities. Majority of the respondents chose to deliver in public health facilities in the index pregnancies.

The far location of health facilities with maternity services which was considered a major constraint by the respondents in this study is a common problem in resource-poor settings. This is supported by previous studies in sub-Saharan Africa and Asia.^7,8,10,14^ With only 60.2% of the respondents who had ever attended an antenatal care visit during their last pregnancy, there is an under-utilization of antenatal care services in this environment. This is essentially similar to the total Nigerian national antenatal prevalence but lower than 94% reported in Zambia.^3,16^

The 56.17% of deliveries carried out by unskilled personnel in this study is slightly lower than the Nigeria national figure of 62% but higher than the reports from the Bahi District, central Tanzania, Eastern rural Nepal and Mukona District of Uganda.^3,7,10,12^ This poses a public health challenge as in spite of 60.2% of antenatal prevalence in the preceding pregnancy of the respondents, it was only 43.8% of the antenatal clinic attendees that delivered in a health facility. The 43.8% of the respondents delivered by skilled birth attendants in this study is marginally higher than the Nigeria national figure of 38%.^3^ This is worrisome as there is no significant progress made towards improving the proportion of pregnant women delivered by the skilled birth attendants in this environment. Some of the reasons given by respondents for the choice of place of delivery such as nearness of the health facility to the respondents, familiarity of the health workers to the respondents, cost of the delivery, improved services, sudden onset of labour, means of transportation and the person who takes decision on the delivery site are essentially similar to the previous reports in Malawi and Uganda.^11,12^

The study showed that more than half of the respondents would prefer to deliver in a public health facility in this index Pregnancy. The factors that influenced their choice of public health facilities were nearness of the public health facilities to the women, familiarity of health workers to the patients, skilled man power, cheaper cost of delivery and improved services. However, rudeness of health workers in public hospitals could be an inhibitory factor that may prevent patients from seeking delivery in public hospitals. A significant proportion of the respondents would also like to deliver in private health facilities if not because of high cost of delivery services, preponderance of quacks and lack of drugs and other services in such facilities. The perceived difference in the quality of care offered by the public andprivate health facilities in this study is similar to the previous reports in Mukono District, Uganda, Kano, Nigeria and rural Orissa, India respectively.^12,15,18^ The perception among the respondents that health workers in public health facilities were more skilled than those in private health facilities in this study is similar to the previous studies in Uganda, Kano and India respectively.^12,17,18^ While it was noticed that health workers in the public health facilities were rude, those in the private facilities were more caring and sympathetic and this was essentially similar to previous reports.^12,19,20^ Contrary to previous reports in Kano,Nigeria,^12^ drugs were more available in public health facilities than private facilities in this study.

This study was limited by the information provided by the respondents which was prone to recall bias. It was also a hospital-based study in which its findings may not be a true reflection in the society.

In conclusion, this study has shown that greater proportion of women will prefer to deliver in a health facility for proper care and attention. However there are barriers militating against the utilization of these facilities for delivery in spite of a significant proportion of the women having attended ante natal clinic. Some of these barriers can be attributed to poverty, inadequate social amenities, the attitude of health workers, illiteracy and influence of family members. This underscores the need for education and women empowerment. The health policy makers should design programmes aimed at ensuring free antenatal and essential obstetric care services in Nigeria. The Federal and State ministries of Health in Nigeria should carry out effective monitoring of health facilities to remove quackery and ensure that such facilities have maintained at least a minimum standard before they are allowed to operate. There is also need for training and re-training of health workers on public relations which will help reduce their rudeness on patients. Government and other stakeholders may provide incentives as one of the ways of enhancing proper utilization of the healthy facilities by these women to ensure safe delivery.

## Acknowledgement

We wish to thank Mr. Benjamin Okorie Ajah,a doctoral student and lecturer in the Department of Sociology and Anthropology,University of Nigeria,Nsukka, for thorough English edition of this manuscript.

## References

1. Millennium development goals. http://www.unicef.org/mdg/maternal/html. Accessed 3rd march 2014.

2. Abouzahr C, Wardlaw T. Maternal mortality in 2000: Estimates developed by WHO, UNICEF and UNFPA. http://www.chidinfo.org/maternal mortality in 2000.

3. National Population Commission (NPC) [Nigeria] and ICF International. 2014. Nigeria Demographic and Health Survey 2013. Abuja, Nigeria, and Rockville, Maryland, USA: NPC and ICF International

4. Gabrysch S, Campbell OM. Still too far to walk: Literature review of the determinants of delivery service use. BMC Pregnancy and Childbirth. 2009;9:34. doi:10.1186/1471-2393-9-34.

5. Gabrysch S, Cousens S, Cox J, Campbell OMR (2011) The Influence of Distance and Level of Care on Delivery Place in Rural Zambia: A Study of Linked National Data in a Geographic Information System. PLOS Medicine 8(1): e1000394. https://doi.org/10.1371/journal.pmed.1000394

6. Wanjira C, Mwangi M, Mathenge E, Mbugua G, Ng’ang’a Z. Delivery practices and associated factors among mothers seeking child welfare services in selected health facilities in nyandarua south district, Kenya. BMC Public Health, 2011; 11: 360.

7. Lwelamira J, Safari J. choice of place of childbirth: prevalence and determinants of health facility delivery among women in Bahi district, central Tanzania. Asian J Sci. 2012; 4(3): 105–112.

8. Pardeshi GS, Dalvi SS, Pergulwar CR, Gite RN, Wanje SD. Trends in choosing place of delivery and assistance during delivery in Nanded district, Maharashtra, India. J Health Popul Nutr. 2011;. 29(1): 71–76.

9. World Health Organisation. Global Health Observatory (GHO) data. www.who.int/gho/urban-health/service/antenatal-care. Accessed march 2015.

10. Ramesh KD. Factors influencing the choice of place of delivery among women in Eastern Rural Napal. Int J maternal and child health. 2013; 30–37.

11. Seljeskog L, Sundby J and Chimango J. Factors influencing women’s choice of place of delivery in Rural Malawi - an explorative study. Afr J Reprod Health 2006; 10(3):66–75.

12. Kkonde, Anthony. Factors that influence pregnant women's choice of delivery site in Mukono district, Uganda, University of South Africa, Pretoria, 2010 http://hdl.handle.net/10500/3601

13. Department of Health Service (DOHS). Kathmandu: Ministry of Health and Population (MOHP) and DOHS. Annual Report 2009/10.

14. Ambreen N, Nishat Z. Women preference for place of delivery; a study at tertiary care hospital. Isra med J. 2013; 5(1).

15. Charan J, Biswas T. How to Calculate Sample Size for Different Study Designs in Medical Research? Indian J Psychol Med.2013;35(2):121–126.

16. Kyei NN, Chansa C, Gabrysch S. Quality of antenatal care in Zambia: a national assessment. BMC Pregnancy Childbirth. 2012 Dec 13;12:151. doi: 10.1186/1471-2393-12-151.

17. Adam YM, Salihu HM. Barriers to the use of antenatal and obstetric care services in rural kano. Niger J Obstet gyneacol. 2002;22 (6):600–603.

18. Alastair A, Pepper K. patterns of health service utilization and perceptions of needs and services in rural orrissa. Health popul planning 2005;20:76–184.

19. Bhatia J, Cleland J. Health care of female out patients in south central India: comparing public and private sector provision. Health popul planning 2004; 19(6):402–409.

20. Russell S. Treatment-seeking behavior in urban sri Lanka: trusting the state, trusting private providers. Soc sci med. 2005; 61: 1396–1407.

